# The Vaginal Microbiome as a Reservoir for Uropathogens: Population-Scale Metagenomics of Women Recently Experiencing BV & UTI

**DOI:** 10.1101/2025.11.27.690987

**Authors:** Sonia N. Whang, Xinyue Wang, Krystal J. Thomas-White, Genevieve Olmschenk, John E. Garza, Pita Navarro, Nicole M. Gilbert

**Affiliations:** Department of Pediatrics, Washington University School of Medicine in St. Louis, MO 63110, USA; Institute for Informatics, Data Science and Biostatistics, Washington University School of Medicine in St. Louis, MO 63110, USA; Evvy, New York, NY 10001, USA; McDonnell Genome Institute, Washington University School of Medicine in St. Louis, MO 63310, USA; Department of Obstetrics and Gynecology, Washington University School of Medicine in St. Louis, MO 63110, USA; Department of Molecular Microbiology, Washington University School of Medicine in St. Louis, MO 63110, USA; Center for Women’s Infectious Disease Research, Washington University School of Medicine in St. Louis, MO 63110, USA

## Abstract

The vaginal microbiome (VMB) influences susceptibility to urogenital infections, yet large-scale, population-based species-level metagenomic studies are rare. We analyzed cross-sectional shotgun metagenomic profiles and linked clinical metadata from 10,003 women across the United States who self-reported recent bacterial vaginosis (BV), urinary tract infection (UTI), both, or neither. Women reporting recent BV or UTI displayed distinct community structures, including higher prevalence of VALENCIA CST-IV subtypes and significantly elevated alpha diversity compared with women who reported no prior diagnosis. Species-level *Gardnerella* profiling revealed that multiple *Gardnerella* species were enriched in the recent BV group but did not differ significantly between UTI and non-UTI groups, refining prior mechanistic hypotheses. Uropathogens such as *E. coli, E. faecalis*, and *S. saprophyticus* were detectable at higher prevalence and relative abundance in women who recently experienced UTI, including among participants who reported recent antibiotic use, consistent with the possibility of residual or recurrent vaginal colonization. These findings demonstrate that microbial signatures associated with recent BV and UTI remain detectable at population scale, provide a high-resolution reference for human vaginal metagenomics, and offer new directions for prevention strategies that consider the vaginal reservoir in recurrent urogenital infections.

## INTRODUCTION

The vaginal microbiome (VMB) is a dynamic and complex microbial ecosystem that plays a central role in maintaining mucosal homeostasis, pathogen defense, and reproductive health. The VMB is classified into community state types (CSTs) based on dominant bacterial taxa ^1^. CST I, II, III, and V are dominated by the *Lactobacillus crispatus*, *L. gasseri*, *L. iners*, and *L. jensenii,* respectively. In contrast, CST IV is characterized by a diverse, polymicrobial community depleted of *Lactobacillus*. The CST IV classification encompasses diverse non-*Lactobacillus* communities but lacks the resolution to distinguish among different dysbiotic states. The VAginaL community state typE Nearest CentroId clAssifier (VALENCIA) classification subdivides CST-IV into specific profiles dominated by *Gardnerella* (IV-A, IV-B), *Prevotella* (IV-C0), or other anaerobes (IV-C1/2/3/4) ^2^. A polymicrobial VMB can be clinically diagnosed as bacterial vaginosis (BV). BV is the most common cause of vaginal discharge among reproductive-aged women ^3^. It is diagnosed by the presence of at least three of four Amsel criteria: (1) thin, homogeneous, grayish-white vaginal discharge; (2) elevated vaginal pH (>4.5); (3) positive whiff test (fishy odor upon addition of potassium hydroxide); and (4) clue cells (epithelial cells coated in bacteria) on microscopy ^4^. Molecular PCR diagnostics like the BD MAX™ Vaginal Panel that detect BV-associated taxa such as *Gardnerella, Atopobium vaginae*, and *Megasphaera* are also available. BV has been associated with serious health risks, such as increased susceptibility to sexually transmitted infections (STIs), pelvic inflammatory disease, urinary tract infections (UTI) and adverse pregnancy outcomes including preterm birth ^5,6^. Globally, the prevalence of BV is high, ranging from 23-29% ^3^. BV is highly recurrent and notoriously difficult to cure; 50%-80% of women will experience a recurrence within 6-12 months ^7,8^. Several intrinsic and sociodemographic factors, including age, race and ethnicity, menopause status, and body mass index (BMI) are associated with increased rates of BV. Black women are more than twice as likely to develop BV compared to white women of European ancestry (non-Hispanic) and show the highest prevalence overall, followed by Hispanic women ^3^. International data consistently show the highest BV rates among younger women in their reproductive years (aged 20-40) and decreasing rates as women age through perimenopause and post-menopause ^9–11^. A meta-analysis estimated an average BV prevalence of approximately 16.9% in postmenopausal women ^12^. Interestingly, the VMB often shifts to lower levels of *Lactobacillus* during perimenopause and post-menopause, despite lower rates of BV diagnoses ^9^, again showing that a polymicrobial VMB is not always definitive of BV or symptoms. A relationship between body mass index (BMI) and BV has been noted ^13^, but this observation is not consistent in all studies ^13–15^. Collectively, studies to date highlight a complex, context-dependent relationship between the VMB and symptomatic BV, underscoring the need to better understand microbial and host factors that predispose to persistent or relapsing disease.

Urinary tract infections (UTIs) are among the most common bacterial infections acquired in the community and in hospitals, affecting millions of patients annually and resulting in substantial healthcare burden. More than 400 million UTI cases occur each year and the global burden of UTIs is rising ^16^. Annually, the cost of diagnosing and treating UTI reaches billions of dollars globally, with approximately $2 billion per year in the United States ^17^. UTI diagnosis can be made based on a combination of symptoms and a positive urine dip-stick analysis or culture. The infection is commonly present with a constellation of lower urinary symptoms including dysuria (pain or burning during urination), urinary frequency, urgency, suprapubic or lower abdominal discomfort, and cloudy or foul-smelling urine that may occasionally contain hematuria (blood in the urine). Most infections in all populations are caused by the Gram-negative uropathogenic *E. coli*. *E. coli* account for 70–90% of community-acquired UTIs and approximately 50% of nosocomial UTI cases ^18^. Beyond *E. coli*, other frequently isolated uropathogens include *Klebsiella pneumoniae, Pseudomonas aeruginosa, Proteus mirabilis, Streptococcus agalactiae, Enterococcus faecalis*, and *Staphylococcus saprophyticus*. Women experience higher rates of UTI and recurrent UTI (rUTI) than men, and the risk of rUTIs may escalate after menopause ^19,20^. Like BV, UTI is highly recurrent; 30-40% of women will experience a recurrence within 6 months, despite successful antibiotic treatment ^21–23^. The gut and the vagina have long been recognized as potential reservoirs for uropathogens that could seed recurrent infections. Multiple studies, encompassing over 1,100 women, have linked the VMB to UTI risk, finding that women with BV or a *Lactobacillus*-deplete VMB are 2 to 13 times more likely to develop UTIs compared to their BV-free counterparts, even after adjusting for age and pregnancy status ^24–27^. Vaginal interventions that restore *Lactobacillus* dominance ^28,29^, including topical estrogen and probiotic *L. crispatus* ^30^ have shown protective effects, significantly reducing the recurrence of UTIs. Furthermore, the vagina is increasingly recognized as a potential reservoir for uropathogens implicated in UTIs. Its anatomical proximity to the urethra, along with factors such as sexual activity and hygiene practices, could facilitate the transfer of vaginal bacteria into the lower urinary tract. Common taxa such as *Gardnerella, Prevotella, Ureaplasma*, and *Lactobacillus* frequently co-occur in vaginal and urine samples, suggesting anatomical migration ^31,32^. A growing body of evidence supports a close microbial connection between the VMB and the urinary tract. Culture-based and 16S rRNA studies have shown that the microbiomes of vaginal and catheterized urine samples display greater microbial similarity with each other than with the gut ^33^. The correlation is particularly strong in women with acute BV ^34^. Thus, vaginal microbes other than recognized uropathogens may also impact UTI susceptibility and severity. For example, our group has demonstrated in an experimental mouse model that transient exposure to *Gardnerella* can induce urothelial exfoliation and exacerbate UTI severity by promoting uropathogenic *E. coli* (UPEC) persistence or by reactivating latent UPEC reservoirs to cause rUTI and pyelonephritis ^35,36^. Specifically, when we exposed mice to *Gardnerella* in their bladders prior to UPEC inoculation, the mice were more susceptible to persistent high-titer UPEC UTI ^35,37^. When *Gardnerella* was inoculated into mice whose bladders harbored quiescent intracellular reservoirs of UPEC from a previous resolved UTI, *Gardnerella* exposure triggered UPEC egress to cause rUTI ^36,38^. Furthermore, *Gardnerella* strains exhibited clade-dependent differences in bladder pathogenicity in our mouse model ^35,38^, underscoring the need to investigate their distinct relationships with UTI in women. These data establish a biologically plausible link between the vaginal microbiome and UTI susceptibility. However, due to the biological and anatomical differences between human and mouse vaginal mucosa and microbiomes,^39^ whether these mechanistic observations translate to human populations—and whether specific *Gardnerella* species contribute differentially to BV versus UTI—remains unclear.

Shotgun metagenomics offers species-level, and in some cases strain-level, resolution of vaginal communities, enabling precise evaluation of microbial signatures linked to BV, UTI, and other urogenital conditions. Despite its advantages, shotgun metagenomics has predominantly been applied in small cohorts or controlled research studies, limiting population-level insights. Large-scale, human metagenomic datasets have the potential to address this gap by capturing a broader range of life-course, demographic, symptom, and treatment variables that influence microbial ecology and infection risk. Here we report on our academic-industry collaboration analyzing shotgun metagenomic profiles and linked metadata from 10,003 women across the United States who self-reported recent BV, UTI, both infections, or neither. To our knowledge, this cross-sectional study represents the largest shotgun metagenomic analysis of the human vaginal microbiome in relation to BV and UTI in a population-based observational dataset. Participants answered a questionnaire which allowed us to evaluate associations between the VMB, demographic, life course, symptoms, diagnoses, and treatments. This dataset enabled us to (1) evaluate associations between recent BV and UTI history and VMB composition at population scale, including CST and VALENCIA community structure; (2) characterize species-level *Gardnerella* profiles in relation to both BV and UTI to refine mechanistic hypotheses generated from prior animal work; and (3) examine the prevalence and relative abundance of uropathogens in the vagina, including among participants who reported recent antibiotic use, to explore whether microbial signatures remain detectable after treatment. Together, these analyses leverage population-sized, high-resolution metagenomics to deepen understanding of vaginal–urinary microbial relationships and provide a critical reference framework for future mechanistic, clinical, and precision-medicine studies focused on recurrent urogenital infections.

## RESULTS

### Participant Characteristics and Group Designations

This retrospective cross-sectional observational study included 10,003 participants that self-collected and submitted vaginal swabs for microbiome characterization via shotgun metagenomic sequencing between November 2022 and May 2024. The participants in the study were distributed across all major geographic regions of the United States, including the Northeast, South, Midwest, and West. Participants were classified into four groups based on their self-reported infection history in the last 30 days (**Figure 1**): recent BV only (BV; n = 4846), recent UTI only (UTI; n = 1185), both recent BV and UTI within the past 30 days (BV&UTI; n = 1053) and never diagnosed with either BV or UTI (ND; n = 2919). There were significant differences in self-reported race/ethnic background (\*\*\*\**p* < 0.0001), menopause status (\*\*\*\**p* < 0.0001), age (\*\*\*\**p* < 0.0001), and body mass index (BMI) (\*\**p* = 0.0022) between groups (**Table 1, Supplementary Table 1**), which is consistent with prior research linking these characteristics to BV and UTI risk as well as VMB composition. No significant differences were present in self-reported pregnancy status. Hence, our data analysis was adjusted for race/ethnicity, age, menopause, and BMI.

**Figure 1:**
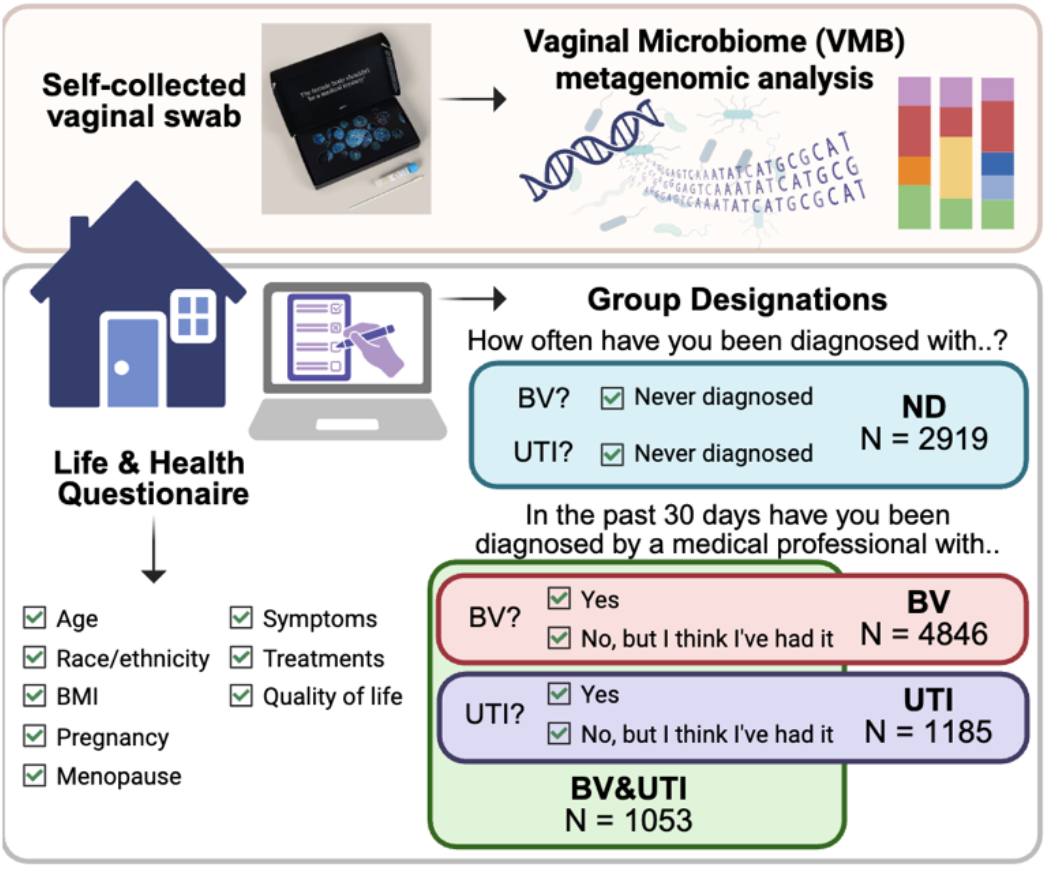
Group designations and study design. Shotgun metagenomic sequencing was performed on self-collected vaginal swabs. The participants completed a health history questionnaire and were classified into study groups based on their answers to the indicated questions.

**Table 1:**
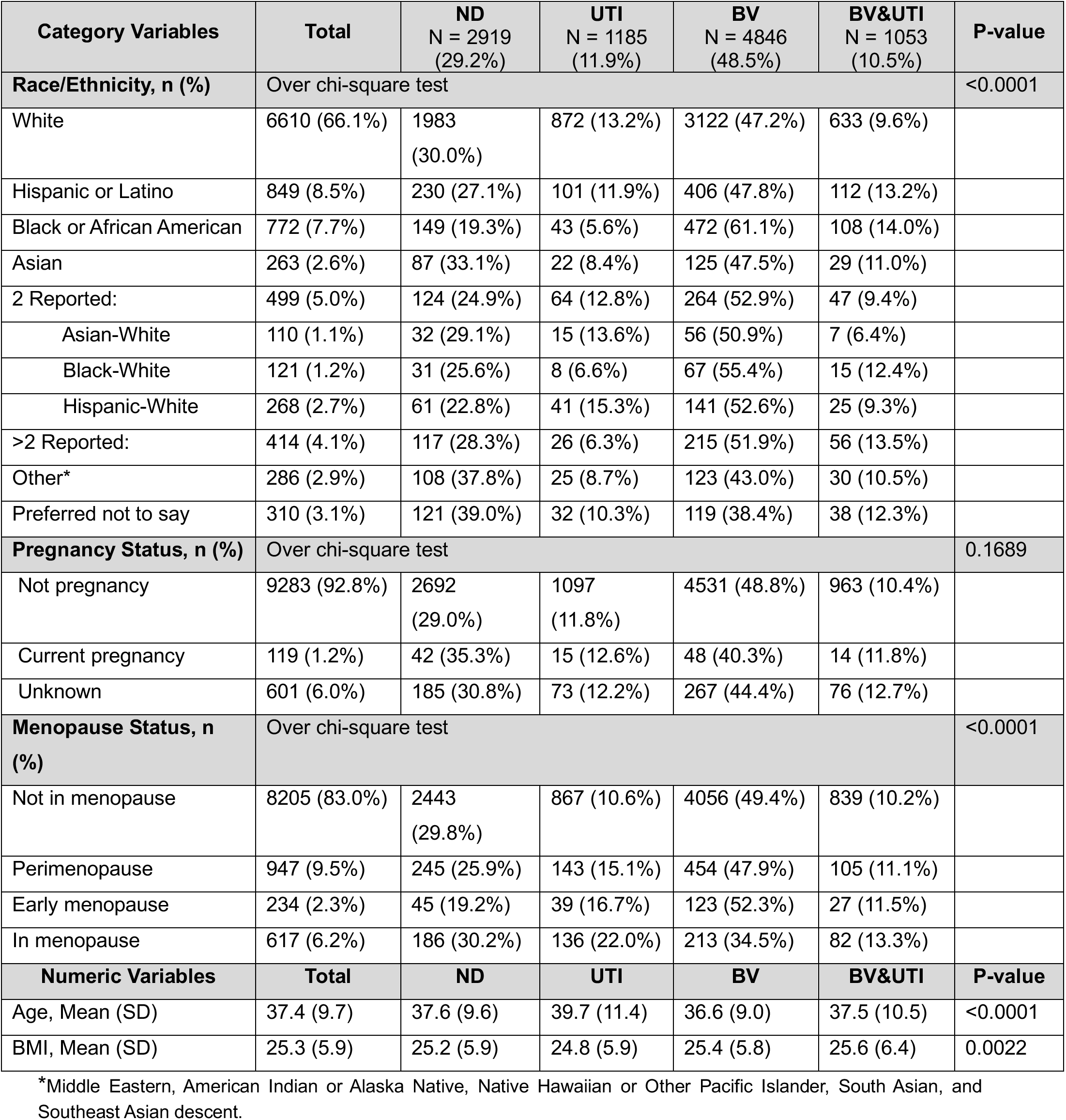
Group Demographics. Categorical variables, two-sided Chi-squared tests. Continuous variables, adjusted LS-mean comparisons and post-hoc multiple comparison tests.

### Community structure differs markedly by recent BV and UTI history

First, we examined the taxa present in the VMB of all the participants included in the study (**Figure 2A-C**). *Lactobacillus* and *Gardnerella* were the most abundant genera. *L. crispatus,* which corresponds to low Shannon diversity CST I and VALENCIA Type I-A /I-B sub-classifications, was the most abundant *Lactobacillus* and VMB dominated by *L. iners* (CST III, VALENCIA IIIA/IIIB), *L. gasseri* (CST II, VALENCIA II), and *L. jensenii* (CST V, VALENCIA V) were also present. The most abundant *Gardnerella* species included *G. swidsinskii* and *G. vaginalis*, followed by *G. piotii*, *G. leopoldii*, and *G. spA*. The distributions of all CST and VALENCIA Types were significantly different between groups (\*\*\*\**p* < 0.0001) (**Supplementary Table 2A-B**). Using a tSNE plot that was generated based on VALENCIA type (**Figure 2B**), clustering of ND samples could be seen in the CST-I region while recent BV samples clustered with CST-IV VALENCIA types (**Figure 2C**). Consistent with established literature ^40^, CST I, III, and IV were the most abundant classifications among all participants (**Figure 2D, Supplementary Table 2A**). The ND and recent UTI groups had a higher proportion of CST I (ND: 41.5%, UTI: 39.2%) and lower CST IV (ND: 29.1%, UTI: 30.7%) compared to both the recent BV and BV&UTI groups (CST I: 24.7%, 26.6%; CST IV: 43.9%, 41.9%, respectively) (**Supplementary Table 2A**). Additionally, the recent BV group was less likely to have CST V (1.8%) and more likely to have CST III (25.1%), while the recent UTI and BV&UTI groups were more likely to have a CST II (6.4%, 5.8%) compared to ND (4.2%) (**Supplementary Table 2A**). To confirm whether these differences in microbial signatures were true in both the women clinically diagnosed and in the women suspecting infections within the last 30 days, we split the infection groups based on whether they answered “yes” <YE> or “I think I’ve had it” <TH>. These subgroups demonstrated similar CST differences when compared to ND, with both the <YE> and <TH> groups having lower CST-I and higher CST-IV (**Supplementary Table 2C**).

**Figure 2:**
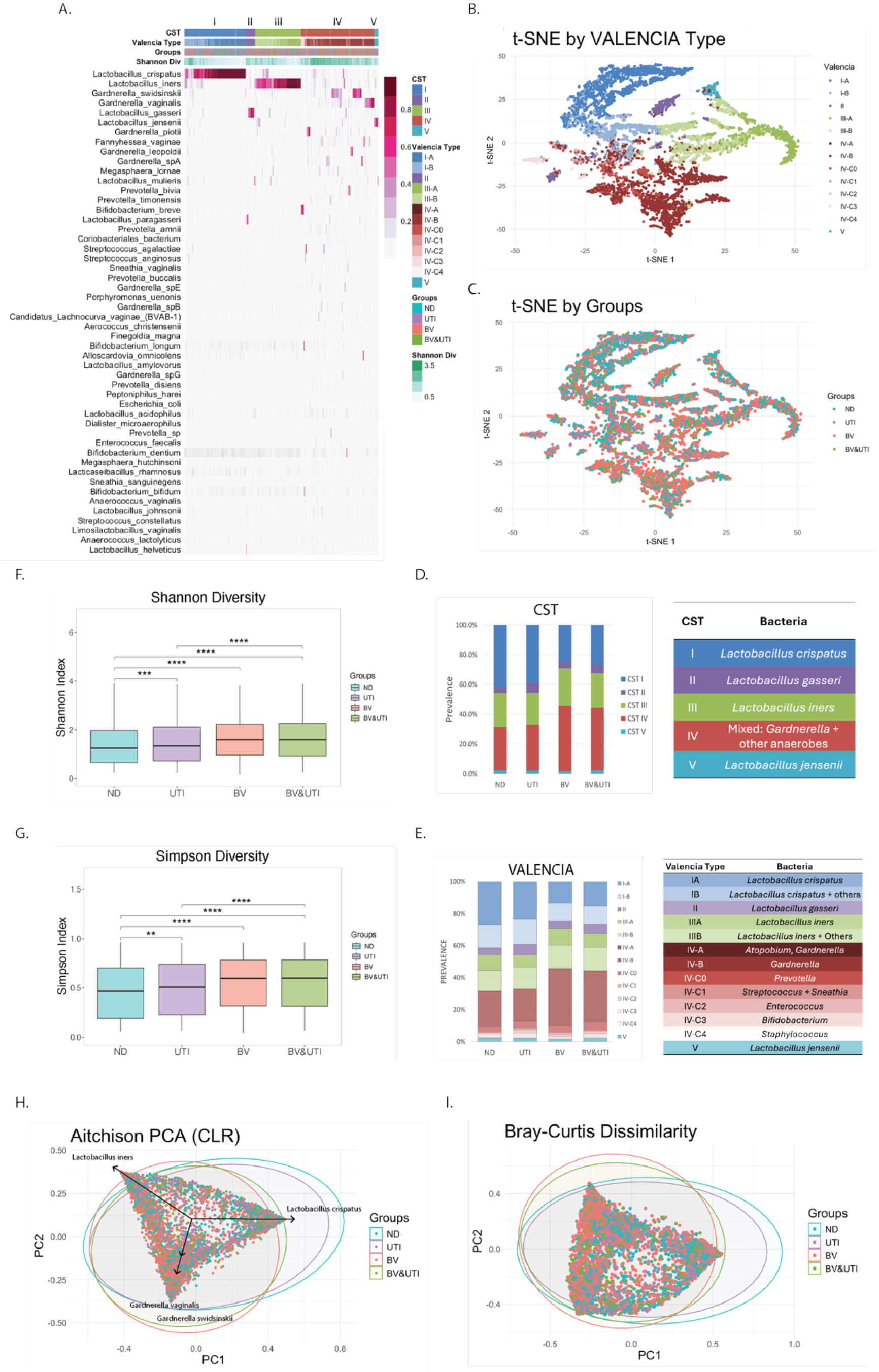
Community structure differs markedly by recent BV and UTI history. **(A)** Heatmaps indicating the CST, VALENCIA type, study group, Shannon diversity, and the relative abundance of the 50 most abundant VMB species from all 10,003 participants. **(B)** *t*-SNE plot of each sample colored by VALENCIA Type. **(C)** *t*-SNE plot from panel B recolored by study group. **(D-E)** Comparative distribution of CST and VALENCIA categories in each study group. Full *p*-values and statistical analysis of the CST and VALENCIA types are listed in **Supplementary Tables S2A-B**, respectively. Vaginal microbiome alpha diversity was assessed using **(F)** Shannon Index and **(G)** Simpson Index and compared using the adjusted LS-mean differences, *p*-values were Bonferroni-adjusted for multiple testing. *\*\*\*p*<0.01, *\*\*\*p*<0.001, *\*\*\*\*p*<0.0001. **(H)** Principal component analysis (PCA) of individual participants, represented by dots and colored based on the groups. Ellipses indicate 95% confidence intervals and bacterial species accounting for the largest differences are indicated based on Aitchison PCA (CLR). **(I)** Principal coordinate analysis (PCoA) of Bray-Curtis distance plot.

While the breakdown of CST between ND and UTI were very similar, the CST-IV VALENCIA Type stratification in the UTI group was more likely to have C0 (4.6%) or C2 (2.1%), which carry an abundance of *Prevotella* and *Enterococcus*, respectively (**Figure 2E; Supplementary Table 2B**). In addition, the BV&UTI group were more likely to classify as CST IV-B (*Gardnerella*/*Atopobium*) (31.6%) and IV-C0 (*Prevotella*) (5.5%), which is consistent with other studies of women with UPEC UTI and BV diagnosis ^41^. In summary, the VMB composition recapitulated known relationships between the VMB and higher CST-IV prevalence in BV, while providing new nuanced differences in the VMB of UTI women.

### Alpha and beta diversity comparisons between groups

Consistent with the polymicrobial nature of the condition, both groups reporting recent BV had significantly higher Shannon & Simpson indexes (alpha-diversity) than ND (\*\*\*\**p* < 0.0001) (**Figure 2F-G**). There was no significant difference in Shannon index noted between recent BV and BV&UTI (*p* = 1.0). Notably, the recent UTI group also exhibited an increase in alpha diversity (Shannon: *** *p* = 0.0002, Simpson: ** *p* = 0.0083) compared to ND. These data demonstrate that women reporting recent UTI have an intrinsically more diverse VMB compared to women in the ND group. We generated a principal component analysis (PCA) based on a Aitchison distance plot (**Figure 2H)** to compare dissimilarity measures and determine the microorganisms that could be attributed to the differences between groups. This analysis identified *L. crispatus* and *L. iners* as the greatest drivers of group variation (44.9%), followed by several *Gardnerella* species (22.6%). The ND group cluster was positioned towards *L. crispatus*, while both the BV and BV&UTI groups were shifted toward *L. iners* and *Gardnerella* species (**Figure 2H**). The UTI group had substantial overlap with ND, with only a slight shift toward the other two groups. A very similar pattern and shift was observed using a PCoA of Bray-Curtis distance plot (**Figure 2I**). Bray-Curtis dissimilarity was significantly different between groups based on an unadjusted permutational multivariate analysis of variance (PERMANOVA) analysis, but was not significant after adjusting for race, age, BMI and menopause status.

### Symptoms and quality of life

Women in the study ranked how much their symptoms affected their quality of life on a scale of 1-10. Women experiencing recent BV or UTI reported higher (meaning worse) scores compared to ND women (\*\*\*\**p* < 0.0001) (**Figure 3A**). Likewise, the rates of reported symptoms were significantly different between the four groups and matched what would be expected for each infection (**Figure 3B**, **Supplementary Table 3**). For example, the women in the recent BV groups were more likely than ND to report that they experienced moderate or severe *excessive discharge* and *odorous discharge* (red cells in **Supplementary Table 3**). Thin vaginal fluid and fishy odor are hallmark and diagnostic features of BV. Moreover, odorous discharge and several vaginal smells (fishy, rotten) were significantly positively associated with BV-associated bacteria, including *Gardnerella, Sneathia,* and *Prevotella,* and were negatively associated with *L. crispatus* (**Figure 3C**). Women in the UTI groups more often experienced the hallmark UTI symptoms of moderate or severe *burning sensation* and *pain while peeing* and also had a higher rate of vulvar pain (purple cells in **Supplementary Table 3**). There were fewer microbial associations with UTI symptoms than there were with BV symptoms.

**Figure 3:**
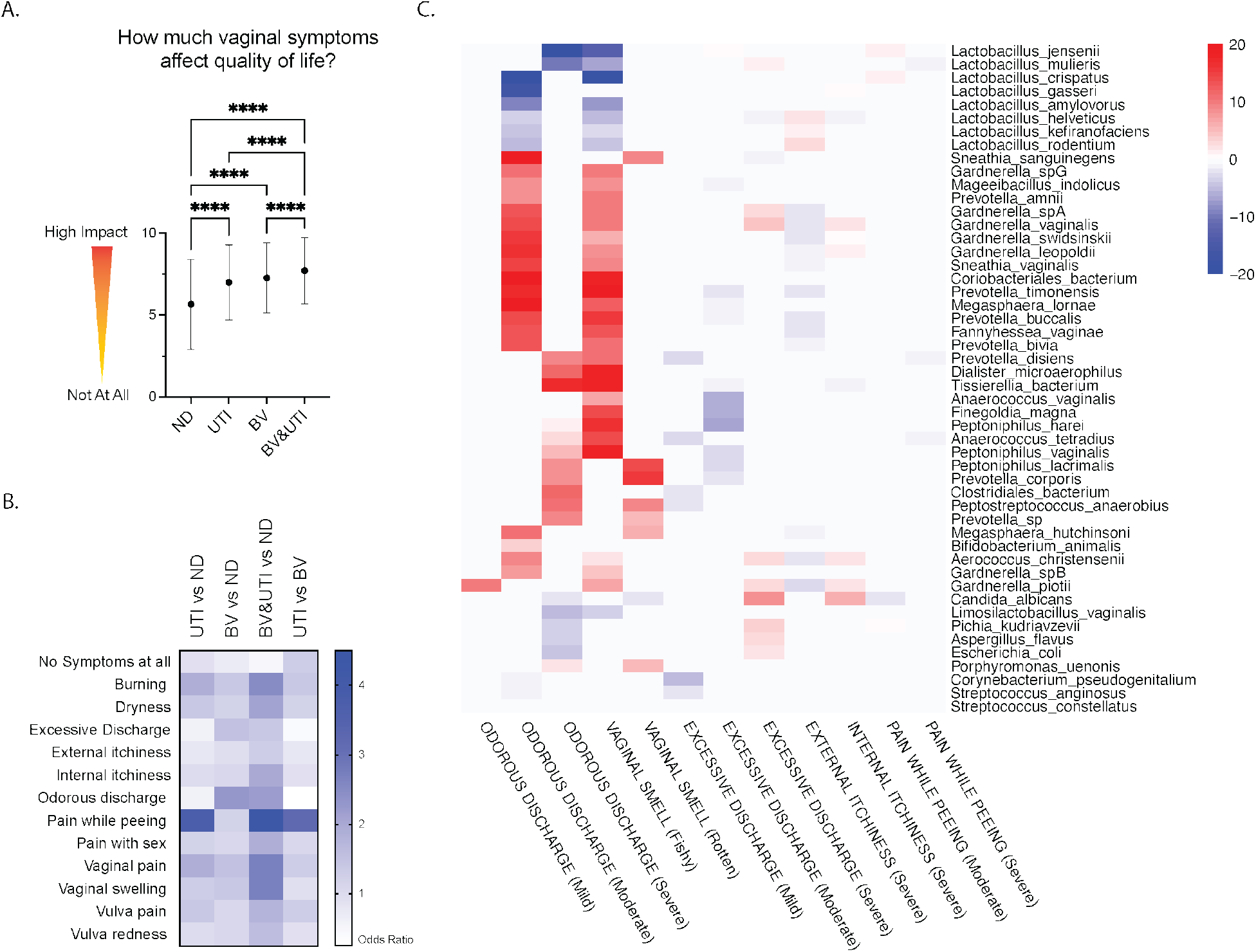
Symptoms and quality of life. **(A)** Self-reported ranking of how much vaginal symptoms affected subjects’ quality of life, analyzed using adjusted LS-mean. *\*\*p*<0.01, *\*\*\*\*p*<0.0001. (**B**) Heatmap of odds ratios comparing self-reported symptoms, derived from Supplementary Table 3. (**C**) Heatmap of MaAsLin2 analysis showing the top 50 significant associations (−log(qval)*sign(coeff)) between bacterial species and common BV and UTI urogenital symptoms.

### Species-level Gardnerella profiles distinguish BV from ND but not UTI from non-UTI

Our previous mouse models demonstrated that urinary tract exposure to *Gardnerella* exacerbates UTI caused by uropathogenic *E. coli* both by promoting persistent infections ^37^ or by triggering recurrence from latent intracellular reservoirs ^38, 36, 35^. We recently reported that only certain *Gardnerella* species can persist in the bladder or cause urothelial damage and inflammation ^42^. We reasoned that the vagina serves as a reservoir for UTI-promoting urinary tract exposures and hypothesized that women experiencing recent UTI might harbor higher levels or distinct species of vaginal *Gardnerella*. Therefore, we compared the relative abundance levels of *Gardnerella* between each group. Approximately 20% of women in each of the recent BV groups (BV: 23.78%, BV&UTI: 20.51%) had a VMB dominated by (>50% relative abundance) *Gardnerella*, which was a significantly higher frequency (\*\*\*\**p* < 0.0001) than in the ND and recent UTI groups (ND: 13.36%; UTI: 8.86%, respectively) (**Figure 4A**). Likewise, both groups reporting recent BV had significantly higher (\*\*\*\**p* < 0.0001) total *Gardnerella* relative abundance (BV: 0.254; BV&UTI: 0.2233) than the ND group (ND: 0.157) (**Figure 4B**). Similar patterns were observed for the most abundant *Gardnerella* species, with significantly higher *G. vaginalis, G. piotii, G. swidsinskii, G. leopoldii, and G. spA* in both recent BV groups relative to the ND group (**Figure 4C-G**). Among the less abundant *Gardnerella, G. spB* (\*\*\*\**p* < 0.0001)*, G. spE* (\*\**p* = 0.0017), and *G. spG* (\*\*\*\**p* < 0.0001) exhibited significantly higher levels in the recent BV compared to the ND group (**Figure 4H; Supplementary Figure 1C and 1E**). *G. spB* was significantly higher in the recent BV compared to the recent BV&UTI group (\*\**p* = 0.0092) (**Figure 4H**), and *G. spD* was significantly higher in the recent BV&UTI compared to the ND group (\**p* = 0.0493) (**Supplementary Figure 1B**). Most samples had multiple *Gardnerella* species and only a small proportion of women in the study had a VMB dominated by a single *Gardnerella* species (**Supplementary Fig 1G-J**). Prior studies have reported both positive and negative correlations between certain *Gardnerella* species ^43, 44^. In our study, pairwise correlation analyses found that the relative abundance levels of all *Gardnerella* species were significantly positively correlated with each other in all four groups (**Figure 4I-L**). The correlations were strongest between *G. vaginalis, G. piotii, G. leopoldii*, and *G. swidsinskii, G. spA, G. spB, and G. spG.* To summarize, our results were consistent with previous studies that have established *Gardnerella* as a BV-associated genera with varying relative abundances between distinct species. Contrary to our hypothesis, there was no apparent association between *Gardnerella* relative abundance or species distribution and recent UTI experience.

**Figure 4:**
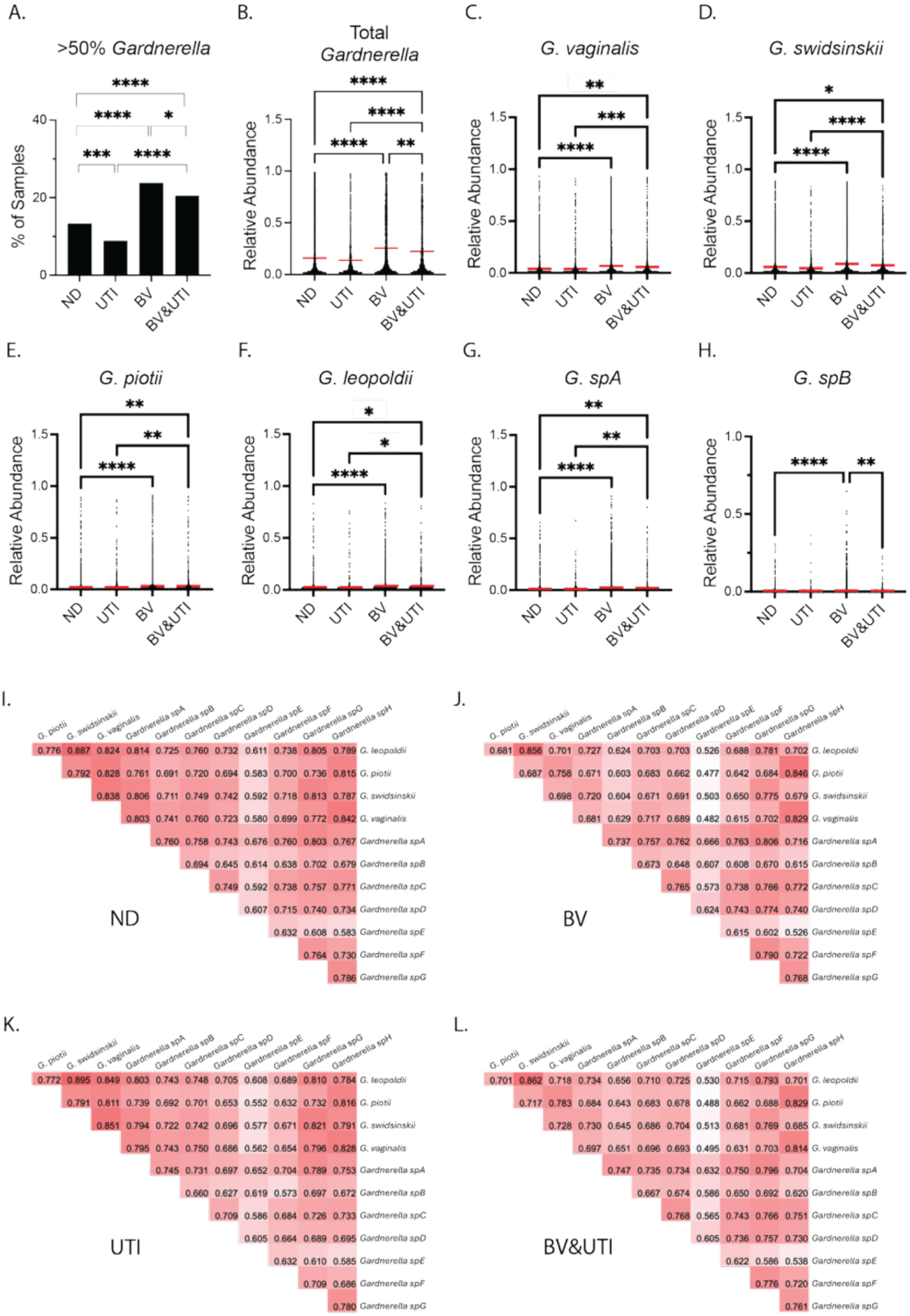
Species-level *Gardnerella* profiles distinguish BV from ND but not UTI from non-UTI. **(A)** Prevalence comparisons of *Gardnerella*-dominant (>50% relative abundance) VMB. \**p*<0.05, \*\**p*<0.01, \*\*\**p*<0.001, \*\*\*\**p*<0.0001, Pearson Chi-square test. Comparisons of species-level *Gardnerella* dominance data are found in Supplementary Figure 1G-J. **B-H** *Gardnerella* relative abundance comparisons using adjusted LS-mean, *p*-values were Bonferroni-adjusted for multiple testing. \**p*<0.05, \*\**p*<0.01, \*\*\**p*<0.001, \*\*\*\**p*<0.0001. Comparative analyses of *Gardnerella spC-spH* relative abundances are provided in Supplementary Figure 1A-F. (**I-L**) Correlation matrices of *Gardnerella* species from each group, using Spearman correlation.

### Vaginal taxa associated with each group are consistent with reported infection history

Having not identified any *Gardnerella* species that were increased in women recently experiencing UTI, next we conducted pairwise linear discriminant analysis (LDA) effect size (LEfSe) analysis to identify which vaginal taxa were enriched in each group. Regarding *Lactobacillus,* there were species-specific associations for in each group. The recent UTI group was enriched for *L. gasseri* relative to the ND group, which was enriched for *L. jensenii* (**Figure 5A**). The ND group had enriched *L. crispatus* and *L. jensenii* relative to the recent BV and BV&UTI groups (**Figure 5B-C**). Women reporting recent BV only were also enriched for *L. iners* (**Figure 5B**). Both groups reporting recent BV were also enriched for several *Gardnerella* species and other BV-associated taxa including *Prevotella*, *Sneathia*, and *Fannyhessea* relative to the ND group (**Figure 5B-C**). Both the recent UTI and BV&UTI groups had enrichment of known uropathogens, including *E. coli*, *K. pneumoniae*, *E. faecalis*, and *S. agalactiae* (**Figure 5A and 5C)**. The recent BV&UTI group was also enriched for *P. mirabilis* and *Candida albicans* (**Supplementary Figure 2C**) compared to ND. Similarly, the recent BV&UTI group could be distinguished from the BV group based on enrichment for *E. coli, E. faecalis, P. mirabilis, K. pneumoniae,* and *C. albicans* (**Figure 5D**). Since LEfSe does not account for confounding factors, we also utilized MaAsLin 2 and included group, race/ethnicity, menopause status, age, and BMI as fixed variables (**Figure 5E**). Again, *L. crispatus* was negatively (blue) associated, whereas bacteria like *Prevotella, Fannyhessea,* and *Sneathia*, were positively (red) associated with both the recent BV and BV&UTI cohorts. Nine *Gardnerella* species were associated with both the recent BV and BV&UTI groups. Among these, *G. piotii*, *G. spA*, and *G. spE* were additionally associated with the recent UTI group, as were the uropathogens *E. coli* and *S. agalactiae* (**Figure 5E**).

**Figure 5:**
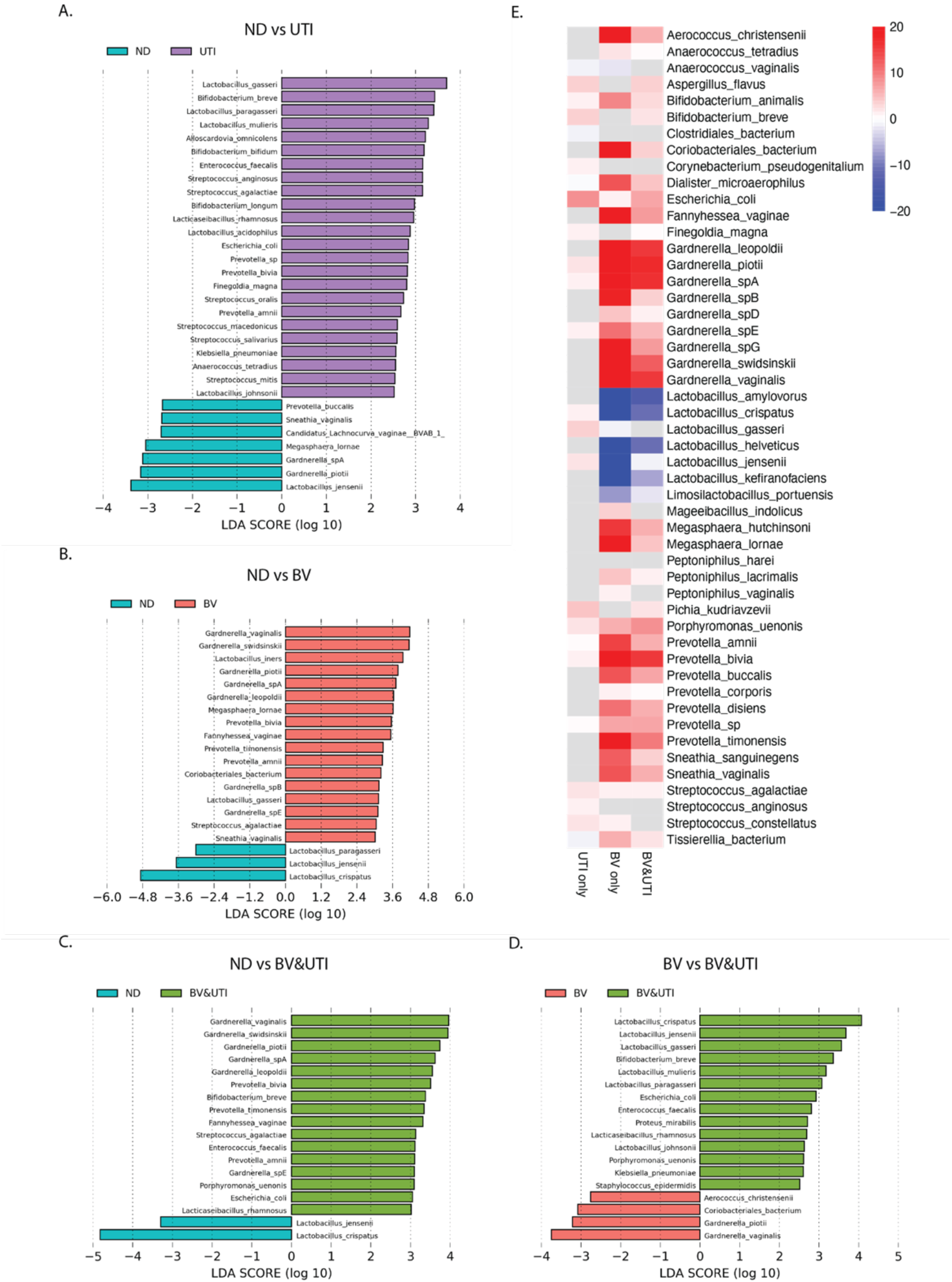
Microbial associations with recent BV or UTI include known BV-associated bacteria and uropathogens. (**A-D**) Pairwise linear discriminant analyses. LDA scores represent the effect size of each abundant species. Species enriched in each group with an LDA score >2.5 are shown for **(A)** ND vs UTI and **(D)** BV vs BV&UTI, and an LDA score > 3.0 is depicted for **(B)** ND vs BV and **(C)** ND vs BV&UTI. For a full list of abundant species with LDA scores > 2.0, see Supplementary Figure 2A-D. **(E)** Heatmap of results from MaAsLin2 analysis showing the top 50 significant associations (−log(qval)*sign(coeff)) between bacterial species and study cohort.

### Vaginal uropathogens are more prevalent and abundant among participants with recent UTI

Results from both the LEfSe and MaAsLin 2 analyses detected associations between vaginal uropathogens and the groups reporting recent UTI. Given these observations, we evaluated the prevalences and relative abundance of the potential uropathogens *E. coli, K. pneumoniae, E. faecalis, P. mirabilis, S. agalactiae, S. saprophyticus, P. aeruginosa,* and *C. albicans* in the VMB collectively and individually. The combined prevalence of these Uropathogens was significantly higher in women who reported recent UTI, even after adjusting for confounding factors (**Table 2**; ND vs. UTI Adj OR 1.4; CI 1.2-1.6; ND vs BV&UTI Adj OR 1.4; CI 1.2-1.6). Individually, *E. coli, E. faecalis, K. pneumoniae*, *S. saprophyticus* and *S. agalactiae* were more prevalent in the VMB of women reporting recent UTI than their ND counterparts. Women with recent BV&UTI exhibited a higher prevalence of *E. coli, E. faecalis,* and *P. mirabilis* compared to the recent BV group; along with these were *K. pneumoniae*, *S. agalactiae* when the recent BV&UTI group was compared to ND. Finally, *E. faecalis* was more prevalent in the recent BV compared to the ND group (**Table 2**).

**Table 2:** Comparison of uropathogen prevalence. Adjusted odds ratio and 95% confidence intervals using the logistic regression model. Presence was defined by a relative abundance of >0.1%, and <0.1% was considered as absence. Significant differences were noted in bold.

| Taxa | UTI vs ND | BV vs ND | BVUTI vs ND | BVUTI vs BV | UTI vs BV | BVUTI vs UTI |
| --- | --- | --- | --- | --- | --- | --- |
| Uropathogens | **1.398**<br>(1.218,1.606) | 1.071<br>(0.977,1.175) | **1.394**<br>(1.207,1.611) | **1.304**<br>(1.137,1.496) | **1.304**<br>(1.144,1.485) | 1.001<br>(0.843,1.189) |
| *Candida albicans* | 1.014<br>(0.812,1.266) | 0.991<br>(0.852,1.154) | 1.029<br>(0.817,1.296) | 1.046<br>(0.841,1.301) | 1.022<br>(0.830,1.259) | 1.033<br>(0.786,1.357) |
| *Enterococcus faecalis* | **2.748**<br>(2.046,3.692) | **1.423**<br>(1.108,1.828) | **2.047**<br>(1.477,2.836) | **1.467**<br>(1.103,1.952) | **1.971**<br>(1.537,2.529) | 0.738<br>(0.533,1.021) |
| *Escherichia coli* | **1.575**<br>(1.356,1.828) | 1.080<br>(0.970,1.203) | **1.420**<br>(1.214,1.662) | **1.319**<br>(1.139,1.528) | **1.456**<br>(1.268,1.671) | 0.907<br>(0.758,1.086) |
| *Klebsiella pneumoniae* | **5.089**<br>(2.911,8.897) | 1.657<br>(0.977,2.810) | **2.648**<br>(1.380,5.078) | 1.585<br>(0.923,2.723) | **3.030**<br>(1.981,4.635) | 0.529<br>(0.299,0.935) |
| *Proteus mirabilis* | 3.387<br>(0.563,20.383) | 1.460<br>(0.282,7.542) | **8.335**<br>(1.673,41.527) | **5.638**<br>(1.708,18.609) | 2.437<br>(0.578,10.270) | 2.304<br>(0.570,9.323) |
| *Pseudomonas aeruginosa* | 0.419<br>(0.122,1.440) | 0.502<br>(0.247,1.021) | 1.110<br>(0.458,2.692) | 2.254<br>(0.906,5.603) | 0.859<br>(0.246,3.003) | 2.619<br>(0.672,10.201) |
| *Staphylococcus saprophyticus* | **2.215**<br>(1.049,4.679) | 1.384<br>(0.752,2.547) | 1.313<br>(0.533,3.237) | 1.572<br>(0.825,2.995) | 1.572<br>(0.825,2.995) | 0.599<br>(0.238,1.513) |
| *Streptococcus agalactiae* | **1.477**<br>(1.236,1.764) | 1.094<br>(0.960,1.245) | **1.232**<br>(1.017,1.493) | 1.127<br>(0.942,1.348) | **1.341**<br>(1.139,1.579) | 0.836<br>(0.673,1.038) |

Uropathogen relative abundances were generally low compared to the levels of *Lactobacillus* and *Gardnerella* species. Notwithstanding, collective uropathogen relative abundance was significantly higher in the recent UTI (\*\**p* = 0.0049) and BV&UTI groups (\*\**p* = 0.0011) compared to ND (**Figure 6A**). *E. faecalis* (\*\*\*\**p* < 0.0001) and *K. pneumoniae* (\**p* = 0.0401) relative abundances were significantly higher in the recent UTI group compared to ND (**Figure 6C and 6E**). The recent BV&UTI group had significantly higher levels of *E. coli* (\*\**p* = 0.0099)*, E. faecalis* (\*\**p* = 0.0035), *K. pneumoniae* (\*\**p* = 0.0028), and *P. mirabilis* (\*\**p* = 0.0097) compared to ND (**Figure 6B-E**). Furthermore, *E. coli* (\**p* = 0.0331), *K. pneumoniae* (\**p* = 0.0241) and *P. mirabilis* (\*\**p* = 0.0046) were significantly more abundant in the recent BV&UTI group compared to BV (**Figure 6B and 6D-E**). We reasoned that the heightened uropathogen abundance might be stronger in the women who were clinically diagnosed with UTI than those who suspected infection. Therefore, we split the groups based on whether the participant answered “yes” <YE> or “I think I’ve had it” <TH>. The presence of higher uropathogen abundance relative to the ND group was exclusive to the <YE> subgroups of recent UTI and BV&UTI (**Supplementary Table 4**). Women that were recently clinically diagnosed with UTI in the last 30 days (UTI <YE>) had significantly more abundant collective uropathogens (\*\*\**p* = 0.0004), and *E. faecalis* (\*\*\*\**p* < 0.0001) and those recently diagnosed with both (BV&UTI <YE,YE>) exhibited higher uropathogen abundance (\*\*\**p* = 0.0004), *E. faecalis* (\**p* = 0.0269) and *K. pneumoniae* (\*\*\*\**p* < 0.0001) in comparison to ND women. This subgrouping also revealed that BV <YE> had significantly higher level of uropathogens (\*\*\**p* = 0.0006), *E. faecalis* (\**p* = 0.0208) and *S. saprophyticus* (\*\**p* = 0.0015) compared to ND. These results are consistent with the clinical association between BV and UTI.

**Figure 6:**
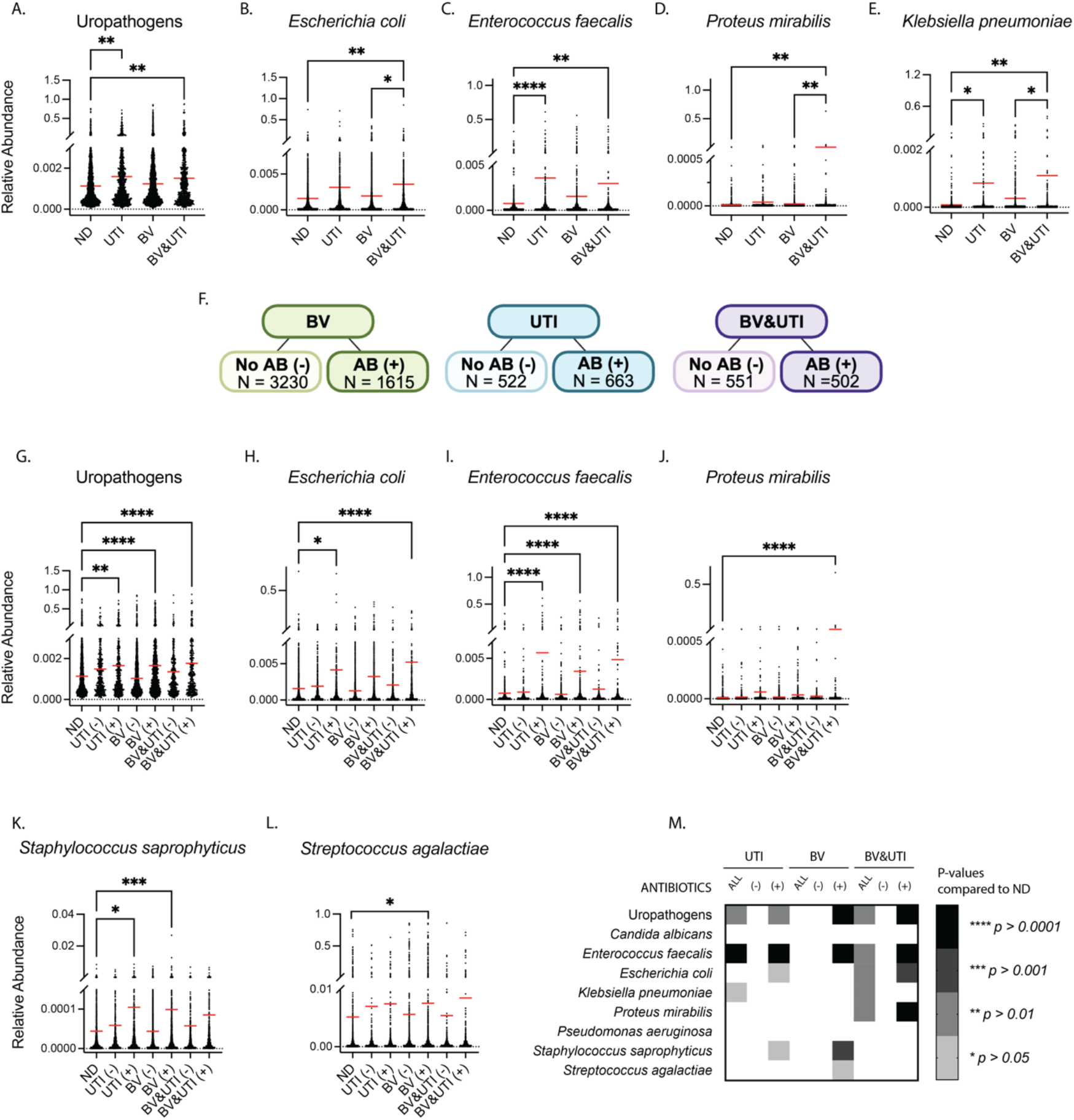
Vaginal uropathogens relative abundances are higher among participants with recent UTI, even those taking antibiotics. **(A-E)** Pairwise comparisons of adjusted uropathogen relative abundances. Statistically significant differences were based on least-squares means, and p-values were Bonferroni-adjusted for multiple testing. Adjusted LS-mean were obtained from general linear models with each group as the primary predictor and age, BMI, race, and menopausal status included as covariates. (**F)** Schematic of sub-analysis stratifying participants within each group based on reported antibiotic use within the last 30 days. Received antibiotic **(AB+)** vs no antibiotics **(AB-)** treatment. **(G-L)** Pairwise comparisons of adjusted relative abundance were based on least-square means, and p-values were Bonferroni-adjusted for multiple testing. **(M)** Heatmap summary of all the significant differences in uropathogen abundance between the AB treated groups **(AB+)**, non-treated group **(AB-)**, and both **(ALL)** compared to ND. \**p*<0.05, \*\**p*<0.01, \*\*\**p*<0.001, \*\*\*\**p*<0.0001.

### Microbial signatures of BV and UTI remain detectable among participants reporting recent antibiotic use

As would be expected, women in the recent BV, UTI and BV&UTI groups were much more likely to report having taken antibiotics in the past 30 days. We reasoned that the patterns of uropathogen colonization could be different between women who did or did not report taking antibiotics (AB) and anticipated that women taking AB might have lower levels of uropathogens, whereas those that did not take antibiotics might have persistent colonization. Therefore, we divided each infection group into participants that either reported using antibiotics (AB+) or not (AB-) (**Figure 6F**). Each infection subgroup was compared to the whole ND group (N = 2919) because there was no difference in uropathogen abundance between the (AB-) and (AB+) ND subgroups (**Supplementary Table 6A**). Contrary to our expectations, there were no significant differences in total or species-level relative abundances between ND and the (AB-) subgroups (**Figure 6G-M** and **Supplementary Table 6A**). However, total Uropathogen relative abundance was significantly higher in the women who took AB across all recent infection groups [UTI (AB+): \*\**p* = 0.0043; BV (AB+): \*\*\*\**p* < 0.0001; BV&UTI (AB+): \*\*\*\**p* < 0.0001] compared to the ND group (**Figure 6G)**. At the level of individual species, the abundance levels were higher than ND for *E. coli* in UTI(AB+) (\**p* = 0.0393) and BV&UTI(AB+) (\*\*\**p* = 0.0009), *E. faecalis* in all three AB subgroups (\*\*\*\**p* < 0.0001), *P. mirabilis* in BV&UTI(AB+)(\*\*\*\**p* < 0.0001), *S. saprophyticus* in BV(AB+) (\*\*\**p* = 0.0007) and UTI(AB+) (\**p* = 0.0397) and *S. agalactiae* in UTI(AB+) (\**p* = 0.0308) (**Figure 6H-M and Supplementary Table 6A**). Since we had observed that heightened uropathogen abundance was specific to the UTI<YE> or BV<YE> subgroups who reported that they had been clinically diagnosed (**Supplementary Table 4**), we examined these subgroups in relation to AB use. This time, some uropathogen species had higher abundances in both the UTI<YE> and UTI<TH> subgroups who reported taking antibiotics (AB+) compared to ND (**Supplemental Table 6B**). In summary, compared to ND, uropathogen abundances were higher in women who reported antibiotic use irrespective of whether the BV or UTI infections they experienced had been clinically diagnosed or were self-suspected.

We observed the opposite pattern to uropathogens when we examined *Gardnerella* abundances with respect to antibiotic use. Although total *Gardnerella* levels were significantly higher in the recent BV groups (\*\*\*\**p* < 0.0001) irrespective of antibiotics, the levels of *G. leopoldii* (\*\*\*\**p* < 0.0001), *G. piotii* (\*\*\*\**p* < 0.0001), *G. vaginalis* (\*\*\*\**p* < 0.0001), *G. swidsinskii* (\*\*\*\**p* < 0.0001), and 5 of the unnamed *Gardnerella* species (*spA, spB, spE, spF, spG*) were only significantly higher in the BV(AB-) (**Supplementary Table 7**). In summary, the relationship between self-reported antibiotic use and relative abundance levels was different between *Gardnerella* and uropathogens.

## Discussion

The importance of the vaginal microbiome in women’s quality-of-life and risk of infection is receiving increased attention outside of academic and clinical settings. The success of recent citizen-science studies of the VMB demonstrates the enthusiasm of women to provide self-collected samples for studies aimed at understanding and improving their health ^45,46^. Women are often looking for answers to urogenital discomfort and recurrent infections that current clinical methodologies and treatments do not resolve. In this large observational study of 10,003 women across the United States, we identified distinct vaginal microbial signatures associated with histories of recent BV, UTI, or both conditions. Using shotgun metagenomics, we found robust enrichment of multiple *Gardnerella* species and anaerobe-dominant VALENCIA CST-IV subtypes among individuals reporting recent BV or BV&UTI, along with significantly elevated alpha diversity relative to participants who had never been diagnosed with either condition. In contrast, individuals reporting recent UTI without BV displayed more heterogeneous community structures and increases in uropathogen prevalence and abundance, highlighting the complexity of vaginal microbiome urinary tract interactions in human populations. Symptom data provided additional context for understanding microbial states. Symptoms typical of BV (odor, discharge, irritation) tracked closely with CST-IV and *Gardnerella*-enriched communities. Relationships that are evident in the literature between the VMB, symptoms, demographics, and BV and UTI were largely recapitulated in this population-based cross-sectional study in which self-sampling and reporting of health parameters occurred in the privacy of the home.

Species-level *Gardnerella* resolution clarified and extended prior observations. Multiple *Gardnerella* species were strongly enriched in women reporting recent BV, but none distinguished participants with and without recent UTI. These results refine hypotheses derived from animal models showing that *Gardnerella* exposure can promote UTI by uropathogenic *E. coli*. Our findings suggest that while *Gardnerella* abundance is a clear biomarker of BV, it may not differentiate UTI risk at the population level, or that such associations may be obscured by heterogeneous infection timing, antibiotic exposures, or species or strain differences not captured in this dataset. Prior studies have implicated *Gardnerella* in BV treatment failure, linking persistence to recurrent BV ^47^. Here we observed a different pattern of *Gardnerella* species between women who did or did not report antibiotic use, but the total *Gardnerella* levels were not significantly different. These associations highlight the need for further study of specific *Gardnerella* clades to understand the clinical significance of species-level dynamics. For example, shotgun metagenomics facilitates the detection of mobile genetic elements, antimicrobial resistance genes, and metabolic pathways, offering deeper insights into the functional potential of vaginal microbial communities. Functional profiles of the metagenomic data could further elucidate the mechanism of infection and recurrence of the microorganisms detected. Moreover, metagenome-assembled genomes could be generated to identify *Gardnerella* genes or pathways associated with BV or UTI history or specific symptoms that women report experiencing.

We also observed that established uropathogens—including *E. coli*, *K. pneumoniae*, *E. faecalis*, and *S. agalactiae*—were detectable at higher prevalence and relative abundance in the VMB among participants reporting recent UTI compared to women never diagnosed with UTI. Although the levels were lower than more typical VMB members, the large sample size equipped us to detect significant associations that would likely be missed in smaller studies. Prior works suggests that the relative abundance levels of uropathogens need not be high to confer risk for UTI. Patients with uropathogenic *E. coli* in their GI tract had increased risk of UTI, even though it only accounted for <1% relative abundance of the GI microbiome ^48^. Notably, uropathogens were detectable in our study among individuals who reported recent antibiotic use. Since information was not collected about the precise infection and antibiotic timing, or whether the antibiotics were taken to treat BV, UTI, or another infection, we cannot conclude that these taxa were detected because they persisted after treatment. However, the higher abundance and prevalence of uropathogens in women experiencing UTI is consistent with the possibility of residual or recurrent colonization and supports the concept of the vagina as a reservoir for uropathogens in women suffering from recurrent UTI. Current clinical practice does not routinely consider the VMB when diagnosing and treating UTIs. Together with prior work, our observational study underscores the need for future longitudinal studies to firmly establish the effects of antibiotics on uropathogen carriage and to consider treatments directed toward eradicating the uropathogen and potential UTI co-conspirators from the vaginal reservoir in future personalized medicine approaches to prevent recurrent UTI.

We acknowledge that there are limitations inherent in this human vaginal metagenomic data study, including the self-collection of vaginal specimens and the self-reporting of infections, symptoms, and treatments. Even though standardized collection kits were used, there could be variation in collection methods by different participants. The infection groups in this study included samples from women who reported that they *experienced* BV or UTI (including those who hadn’t been diagnosed but believed they had it) sometime in the past 30 days. We do not have specific dates of when the infections occurred, and diagnoses were not confirmed by a clinician at the time of sample collection. Approximately 85% of the women who answered “I think so” also reported that they had been diagnosed with the same infection in their lifetime (BV 84.2% = 3425/4066; UTI 85.3% = 1031/1209) and were therefore equipped to recognize their symptoms associated with past clinical diagnoses. We note that it is common practice for clinicians treating patients with a history of rBV or rUTI to prescribe antibiotics based on symptom onset without a clinic visit. Antibiotic use was reported in the last 30 days and should not have occurred during the 7 days immediately prior to sample collection (based on the standard submission instructions to avoid antibiotics for one week), but we do not know the exact timing or type of antibiotics used. The ND group in this study included women who reported that they had never been diagnosed with BV or UTI in the past nor did they think they had it within the last 30 days, but our study design did not consider the reporting of other infections as exclusion criteria. Therefore, although nearly 80% (2333/2919) of the ND group reported no other urogenital infections in the last 30 days, this group was not devoid of infections and should not be considered the same as a “healthy” control. Nearly all women in the study reported that they had been diagnosed with at least one urogenital infection in their lifetime. Despite these limitations, the VMB composition, symptoms, and demographic patterns in each group in this study matched strikingly well with what has been reported from clinical studies and set the stage for future functional analyses such as gene/KEGG enrichment with this dataset that will expand our knowledge of microbial ecology in the vagina.

Together, these findings advance understanding of vaginal microbial ecology at population scale and highlight several implications for clinical practice and future research. First, the strong enrichment of *Gardnerella* and other BV-associated species in women reporting recent BV despite recent antibiotic use supports the current idea that microbial persistence drives recurrent BV. Second, the detection of uropathogens despite recent antibiotic use underscores the need for improved therapeutic strategies that address vaginal reservoirs, promote sustainable *Lactobacillus* colonization, and disrupt microbial communities associated with recurrent UTI. Third, the overlapping symptoms and microbial signatures in women recently experiencing both BV and UTI in close succession supports integrated approaches to managing these infections, rather than treating them as isolated conditions. Fourth, this study demonstrates the feasibility and value of leveraging human vaginal metagenomic data, paired with detailed symptom, treatment, and metadata, for translational insights that complement traditional clinic-based studies. Here we focused on BV and UTI because of our basic science research expertise. We note that we also observed a positive association between *Candida albicans* and *vulvar redness* and *internal itchiness,* which are hallmark symptoms of vulvovaginal *Candidiasis* (VVC). Therefore, our study provides a framework for population-based investigations of other urogenital infections.

## Methods

### Sample collection and questionnaire

All study participants provided informed consent, and study procedures were conducted in accordance with protocols approved by the federally accredited Institutional Review Board of Viome Institutional Review Board (IRB# 20220118.evvy). A sample collection kit (Copan, Murrieta, CA, USA) was shipped directly to the participant who self-collects a vaginal swab and ships the sample at ambient temperature to a CLIA, CAP, and CLEP certified lab (CLIA 45D1086390, CAP 7214171, PFI 9433 Microgen DX, Lubbock, TX, USA). Vaginal samples and metadata were collected between November 2022 and May 2024. The analysis and measurements were taken from distinct vaginal samples. Women filled out an online questionnaire reporting their demographic information. In Table 1, women experiencing perimenopausal symptoms as well as women diagnosed perimenopausal by a doctor were both combined under ‘Perimenopause.’ ‘Early menopause’ included women who experienced hysterectomy, uterine ablation, primary ovarian insufficiency, chemotherapy, etc.

### Group Designations

Inclusion required patients to have completed both the health questionnaire and vaginal microbiome characterization via shotgun metagenomic sequencing through Evvy’s VMB telehealth service platform. Participants were stratified based on their responses to an initial survey at the time of self-collected sample submission (**Figure 1**). Participants were asked, *“<u>In</u> <u>the past 30 days</u>, have you been diagnosed by a medical professional with any of the following infections?”* They could answer *“Yes”*, *“Not diagnosed, but think I’ve had this in the past 30 days”*, or *“No”* to UTI and BV selections (in addition to other urogenital infections not analyzed in this study). Women who responded with *“Yes”* or *“Not diagnosed, but think I’ve had this in the past 30 days”* to either infection were included in the corresponding **BV (**N = 4846) or **UTI** (N = 1185) groups. If they responded positively to the question for both infections, then they were included in the **BV&UTI** (N = 1053) group. Women who answered *“No”* to both infections and also answered *“Never Diagnosed”* to the question *“How often have you been diagnosed with each of the following conditions?”* were placed in the Never Diagnosed (**ND**; N = 2919) group. Participants were instructed to “avoid testing while menstruating, as well as if they have had sex, used an oral antifungal, vaginal suppositories, or vulvar topical cream in the last 24-48 hours, or taken antibiotics within the last 7 days.”

### Sequencing

Samples were processed as previously described ^49^, which includes a chemical and mechanical lysis, host depletion with PMA [15], chemical lysis using metapolyzyme (Sigma, St. Louis, MO, USA), and DNA extraction using an automated extraction handling instrument (KingFisher FLEX, Temecula, CA, USA, Thermo Scientific, Waltham, MA, USA). Each extraction batch includes a negative control (NC, water) and a positive control containing a known mixture of bacteria and fungi. NGS libraries were prepared, multiplexed, quality checked, and sequenced on the NovaSeq 600 (Illumina, San Diego, CA, USA). Each sequencing run included the extraction negative and positive controls, no template controls, and reagent controls. Sequencing data from each participants’ sample were processed through Evvy’s validated pipeline, which is specifically designed to characterize the vaginal microbiome. Samples that passed detection thresholds were reported to the participant ^49^. Participant data collection on the platform included questionnaires documenting symptomatology, relevant clinical diagnoses, and demographic information.

### Metagenomic analysis

Metagenomic data analysis was conducted via Evvy’s previously published and validated bioinformatics platform that includes quality control, host depletion, and taxonomic profiling ^49^. Trimmomatic [16] was used to trim and filter raw reads to remove low quality bases. Following quality filtering, remaining data are further refined by eliminating human DNA sequences by mapping to the latest human reference (GRCh38). Samples are required to have at least 20,000 non-human reads to continue through the bioinformatic pipeline, based on our previously published validation studies ^49^. The reads are then aligned to a proprietary reference database generated from a curated collection of vaginal microbial genomes to generate taxonomic relative abundance profiles^49^. The unique signatures comprising the database are computed from microbial whole genomes isolated from the urogenital tract and contain more than 4000 microbial genomes, representing 700 bacterial species. Between January and March 2023, genomes were collected from publicly available repositories (NCBI and EBI) with any isolate documented to be collected from “urogenital” or “vaginal” sources. To accurately represent the phylogenetic diversity of underrepresented taxa, some genomes were included from other human microbiome sites. Public repositories are incomplete in their representation of the vaginal microbiome; for this reason, we also included metagenome-assembled genomes generated from Evvy test data. *Gardnerella* species were determined as we previously described, using a genome dataset created from publicly available (as of 2023) genomes (N=209) and metagenome assembled genomes generated from Evvy samples (N=280). Species were identified using average nucleotide identity (ANI) of whole genomes with a 95% similarity cutoff ^50^. *G. vaginalis* and *G. piotii* each contained 2 distinct sub-species variants, but these were combined in this study. There were many unclassified *Gardnerella* sp. genomes, but we classified these genomes into subtaxa groups *spA-spH*. Previously established VALENCIA methods ^2^ were used to classify the CSTs and Valencia types for each sample.

### Statistical analysis

Statistical analyses were conducted using SAS Studio (SAS Studio via SAS OnDemand for Academics), running Base SAS 9.4 (M8) and SAS/STAT 15.3 (SAS Institute Inc., Cary, NC); Prism software 10 (version 10.4.1, GraphPad Software Inc, San Diego, CA); Python (version 2.7.12, Python Software Foundation, Delaware, US); and R (version 4.5.0, R Foundation for Statistical Computing, Vienna, Austria). Our primary objectives were to examine whether vaginal microbiome composition differed between groups and whether the relative abundances of *Gardnerella* species and established uropathogen species differed between groups. Clinical subgroups included: Never diagnosed (ND; reference), UTI, BV, and BV&UTI. *P*-values < 0.05 were considered significant. Relative abundance values were derived from metagenomic sequencing data and log-transformed to improve normality where necessary. All models were checked for assumptions of normality, homoscedasticity, and multicollinearity. We used generalized linear models (GLM) to evaluate differences in mean relative abundance of *Gardnerella* and uropathogens across groups. The primary predictor was the clinical group, while age, body mass index (BMI), self-reported race/ethnicity, and menopausal status were included as covariates to control for potential confounding. Least-squares mean (LS-mean) were estimated and pairwise differences between groups were assessed using Bonferroni-adjusted *p*-values and 95% confidence intervals. This approach allowed us to identify statistically significant differences in *Gardnerella* (**Figures 4B-H; Supplementary Figures 1A-F**) and uropathogen (**Figure 6**) abundance while accounting for demographic heterogeneity in the study population.

To assess the association between *Gardnerella* and uropathogen abundance and odds of UTI and/or BV, we conducted multivariable logistic regression analyses. The outcomes were binary clinical variables, and the independent variable of interest was the relative abundance of *Gardnerella* or uropathogenic species. Models were adjusted for age, BMI, race, and menopausal status. Adjusted odds ratios with 95% confidence intervals were reported. In exploratory models, we also examined effect modification by menopausal status and race using interaction terms.

To control for inflated type I error due to multiple comparisons, Bonferroni correction was applied to all pairwise comparisons within the GLM framework. For logistic regression models with multiple microbial predictors, we used false discovery rate correction when appropriate.

To compare categorical microbial abundance classifications (Dominance vs Non-dominant; Presence vs. Absence) across study groups, we performed two-tailed Chi-square tests of independence. We determined *dominance* as 50% microbial relative abundance or higher (**Figure 4A; Supplementary Figures 1G-J**), and *presence* as 0.1% relative abundance or above (**Table 2**) to ensure analytical robustness. These analyses evaluated whether the prevalence of *Gardnerella* species and uropathogens differed among the four clinical subgroups (ND, UTI, BV, and BV&UTI). Pairwise comparisons between study groups (UTI vs ND, BV vs ND, BV&UTI vs ND, UTI vs BV, BV&UTI vs UTI, BV&UTI vs BV) were conducted, and the resulting Chi-square *p*-values were summarized in tables to indicate statistically significant differences in species-level distributions. All categorical analyses were two-sided, and *p* < 0.05 was considered statistically significant.

### Microbiome features, diversity, and association analyses

A heatmap and hierarchical clustering based on Euclidean distance, depicting patterns of abundance from the metagenomic data of each participant was constructed using the ‘pheatmap’ package within R. The heatmap annotations in Figure 2A display the VMB profiles classified based on established community state type (CST) and VALENCIA categories, the Shannon diversity index values, and the study group. For Shannon & Simpson diversity, LS-mean were estimated and pairwise differences between groups were assessed using Bonferroni-adjusted *p*-values and 95% confidence intervals. Principal component analysis (PCA) was used to evaluate the similarities or differences in the composition of the sample communities based on Aitchison distances (Euclidean distance on CLR (centered log-ratio) transformed data) (**Figure 2H**) using the ‘compositions’ package. Principal coordinate analysis (PCoA) based on Bray-Curtis distances (**Figure 2I**) utilized the ‘adonis’ package (<u>79</u>). Alpha diversity (Shannon and Simpson Indexes) and beta diversity (Bray-Curtis Index) were carried out using the packages ‘vegan’ and ‘ggplot’. Significance of Bray-Curtis dissimilarity was analyzed using pairwise permutational multivariate analysis of variance (PERMANOVA) and performed with 999 permutations using the ‘vegan’ package and ‘adonis’ function. The metadata associations with microbial taxa in **Figures 3C** and **5E** were determined using the MaAsLin2 package in RStudio (Figures 3C & 5E; Supplementary Figure 3) with default parameters (normalization = TSS, transformation = LOG, analysis_method = LM, max_significance = 0.25, min_abundance = 0.0 and min_prevalence = 0.1) with the exception that the cohort and demographic associations in **Figure 5E** were performed with min_abundance = 0.0001 and min_prevalence = 0.1). The correlation matrices of *Gardnerella* species (**Figure 4I-L**) were produced by GraphPad Prism analysis, using Spearman correlation and analyzed by a two-tailed test. Differential analysis of taxa was performed using the Python package LEfSe v1.0 (**Figure 5A-D**). In addition, *p* < 0.05 and LDA > 2 obtained by linear discriminant analysis were considered statistically significant (**Supplementary Figure 2A-D**). Data visualization was performed with GraphPad Prism, Python, and RStudio.

## Data availability

The pre-computed relative abundance tables underlying all analyses and figures in this study are available at GitHub under project titled “Evvy-BV-UTI”. Password for Excel files: BVUTImicrobiome https://github.com/GilbertLabWashU/Evvy-BV-UTI. Metagenomic sequencing files are available at the NCBI Sequence Read Archive under BioProject PRJNA1460779 https://dataview.ncbi.nlm.nih.gov/object/PRJNA1460779?reviewer=nntpiee8v5pgvb45nbdj75210v

## Code availability

Analysis code is available at GitHub under project titled “Evvy-BV-UTI” https://github.com/GilbertLabWashU/Evvy-BV-UTI

## Acknowledgements

The authors thank David Lyttle, Rob Markowitz for relative abundance data preparation and assistance with data deposition; Ann Rosen, Dr. Drew Schwartz, Dr. Michael White, Dr. Tychele Turner, and Dr. Christopher Miller for computational and bioinformatics assistance and guidance in processing data and creating figures in R and Python; Dr. Fan Zhang for her biostatistical expertise and initial guidance in the analysis; and Dr. Morgan Timm for proofreading and providing clinical insights and ideas.

## Author contributions

Sonia N. Whang – data preparation and analysis, concept, study design, biostatistical analysis, manuscript draft, figure preparation

Xinyue Wang – biostatistical analysis, table preparation, manuscript draft

Krystal J. Thomas-White – data preparation, concept and expertise, study design, manuscript editing

Genevieve Olmschenk – data preparation and analysis, manuscript editing

John E. Garza – data processing through Python, data figure creation, and data deposition Pita Navarro – concept, study design, manuscript editing

Nicole M. Gilbert – concept and expertise, study design, manuscript draft and editing, resources, oversight

All authors read and approved the manuscript.

## Additional information

## Supplementary information

Supplementary Figure 1: *Gardnerella* less abundant & 50% dominant species

Supplementary Figure 2: Lefse – LDA score > 2.0

Supplementary Table 1: Group Demographics – Odds Ratio (OR)

Supplementary Table 2A: CST – Summary & OR

Supplementary Table 2B: VALENCIA Types – Summary & OR

Supplementary Table 2C: CST <YE vs TH> – Summary & OR

Supplementary Table 3: Symptoms – Summary & OR

Supplementary Table 4: Uropathogen <YE vs TH> – Abundance Difference

Supplementary Table 5: Treatments – Summary & OR

Supplementary Table 6A: Uropathogen Antibiotics – Abundance Difference

Supplementary Table 6B: Uropathogen Antibiotics <YE vs TH> – Abundance Difference

Supplementary Table 7: *Gardnerella* Antibiotics – Abundance Difference

