## Supplemental Figures for "The Vaginal Microbiome as a Reservoir for Uropathogens: Population-Scale Metagenomics of Women Recently Experiencing BV & UTI"

**Supplementary Section**

Supplementary Figure 1A-J: *Gardnerella* less abundant & 50% dominant species

Supplementary Figure 2A-D: Lefse – LDA score > 2.0

(Excel Spreadsheet: Table Captions)

Supplementary Table 1: Group Demographics – Odds Ratio (OR)

Supplementary Table 2A: CST – Summary & OR

Supplementary Table 2B: VALENCIA Types – Summary & OR

Supplementary Table 2C: CST <YE vs TH> – Summary & OR

Supplementary Table 3: Symptoms – Summary & OR

Supplementary Table 4: Uropathogen <YE vs TH> – Abundance Difference

Supplementary Table 5: Treatments – Summary & OR

Supplementary Table 6A: Uropathogen Antibiotics – Abundance Difference

Supplementary Table 6B: Uropathogen Antibiotics <YE vs TH> – Abundance Difference

Supplementary Table 7: *Gardnerella* Antibiotics – Abundance Difference

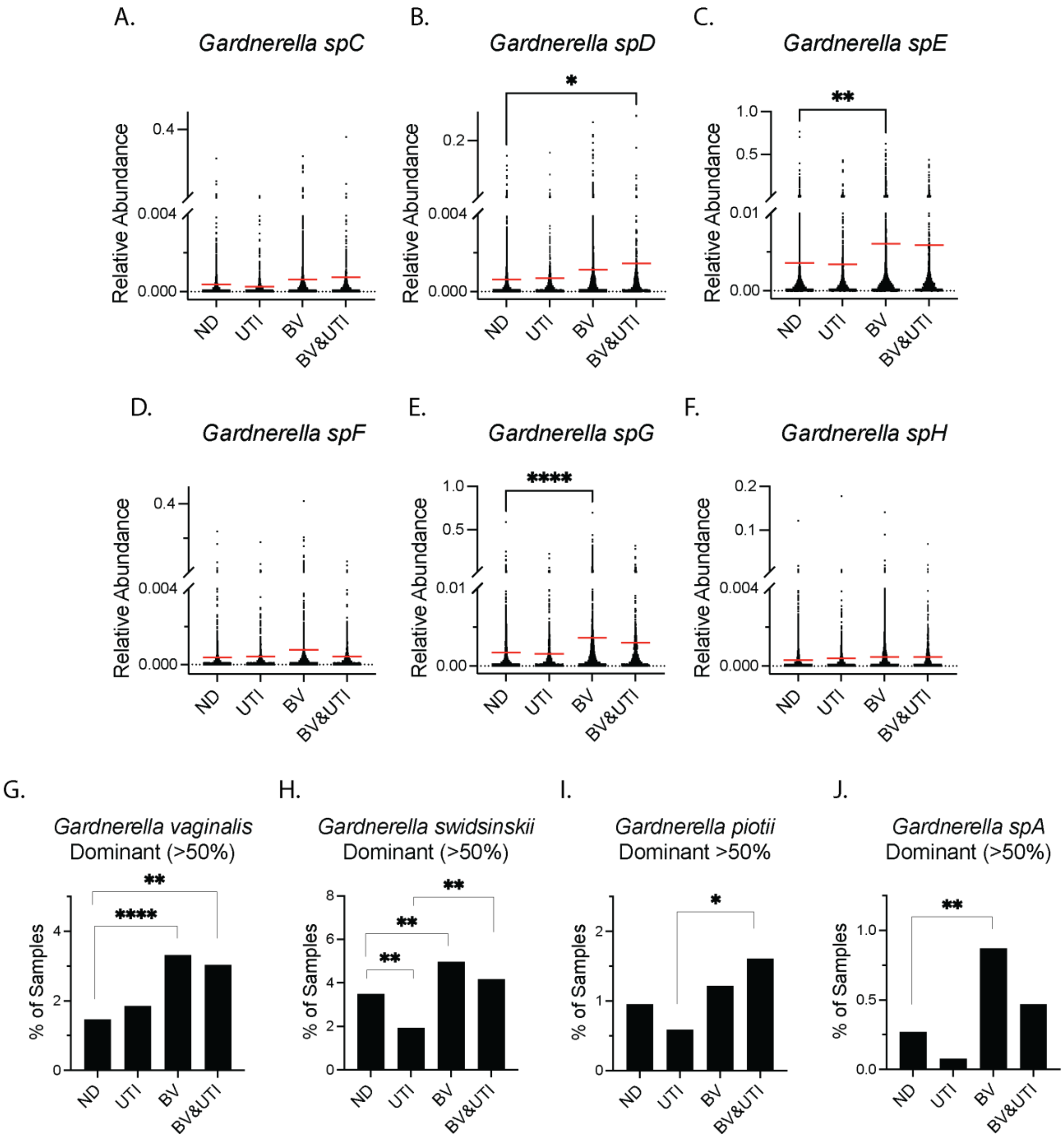


**Supplementary Figure 1**: *Gardnerella*. In panels **A-F**, comparative relative abundance of species-level analysis of *Gardnerella* taxa **(a)** *spC,* **(b)** *spD,* **(c)** *spE,* **(d)** *spF,* **(e)** *spG,* **(f)** *spH*, using adjusted LS-mean, *p*-values were Bonferroni-adjusted for multiple testing. In panels **G-J**, Pearson Chi-square test to compare the prevalence of dominant *Gardnerella* species, defined by a 50% relative abundance or higher, between groups of which significant differences were observed in **(g)** *Gardnerella vaginalis*, **(h)** *Gardnerella swidsinskii*, **(i)** *Gardnerella piotii*, and **(j)** *Gardnerella spA*.

25 **A)**

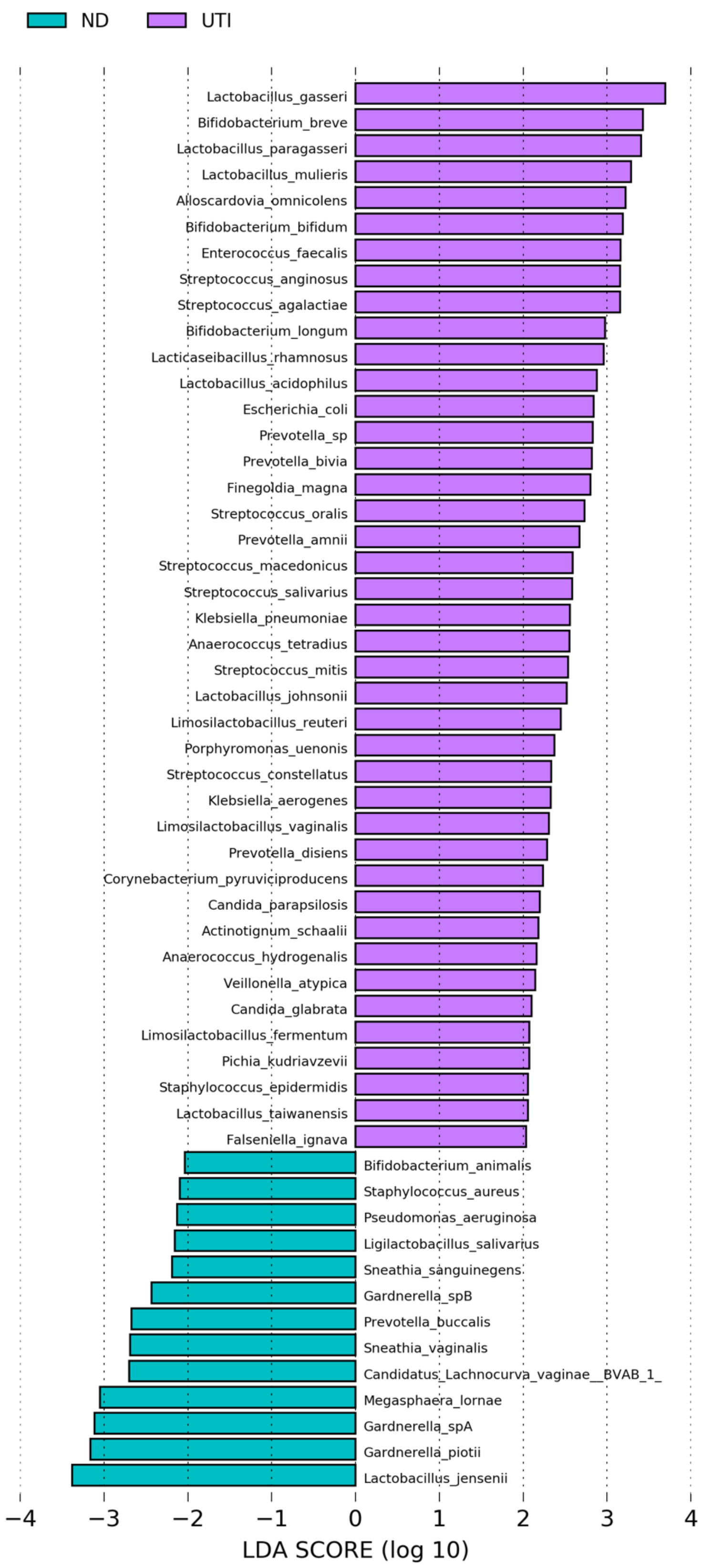

ND
UTI
Lactobacillus_gasseri
Bifidobacterium_breve
Lactobacillus_paragasseri
Lactobacillus_mulieris
Alloscardovia_omnicolens
Bifidobacterium_bifidum
Enterococcus_faecalis
Streptococcus_anginosus
Streptococcus_agalactiae
Bifidobacterium_longum
Lacticaseibacillus_rhamnosus
Lactobacillus_acidophilus
Escherichia_coli
Prevotella_sp
Prevotella_bivia
Finegoldia_magna
Streptococcus_oralis
Prevotella_amnii
Streptococcus_macedonicus
Streptococcus_salivarius
Klebsiella_pneumoniae
Anaerococcus_tetradius
Streptococcus_mitis
Lactobacillus_johnsonii
Limosilactobacillus_reuteri
Porphyromonas_uenonis
Streptococcus_constellatus
Klebsiella_aerogenes
Limosilactobacillus_vaginalis
Prevotella_disiens
Corynebacterium_pyruviciproducens
Candida_parapsilosis
Actinotignum_schaalii
Anaerococcus_hydrogenalis
Veillonella_atypica
Candida_glabrata
Limosilactobacillus_fermentum
Pichia_kudriavzevii
Staphylococcus_epidermidis
Lactobacillus_taiwanensis
Falseniella_ignava
Bifidobacterium_animalis
Staphylococcus_aureus
Pseudomonas_aeruginosa
Ligilactobacillus_salivarius
Sneathia_sanguinegens
Gardnerella_spB
Prevotella_buccalis
Sneathia_vaginalis
Candidatus_Lachnocurva_vaginae__BVAB_1_
Megasphaera_lornae
Gardnerella_spA
Gardnerella_piotii
Lactobacillus_jensenii
−4
−3
−2
−1
0
1
2
3
4
LDA SCORE (log 10)

26 **B)**

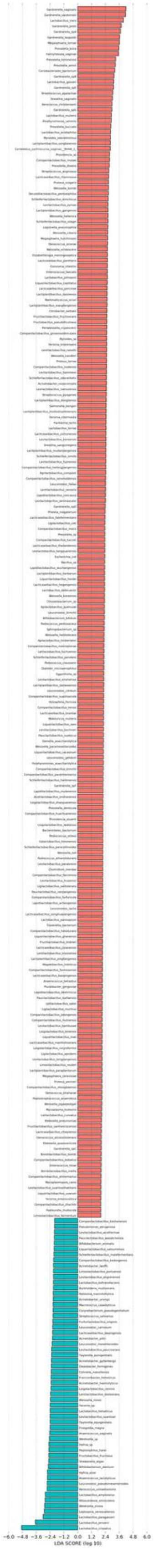

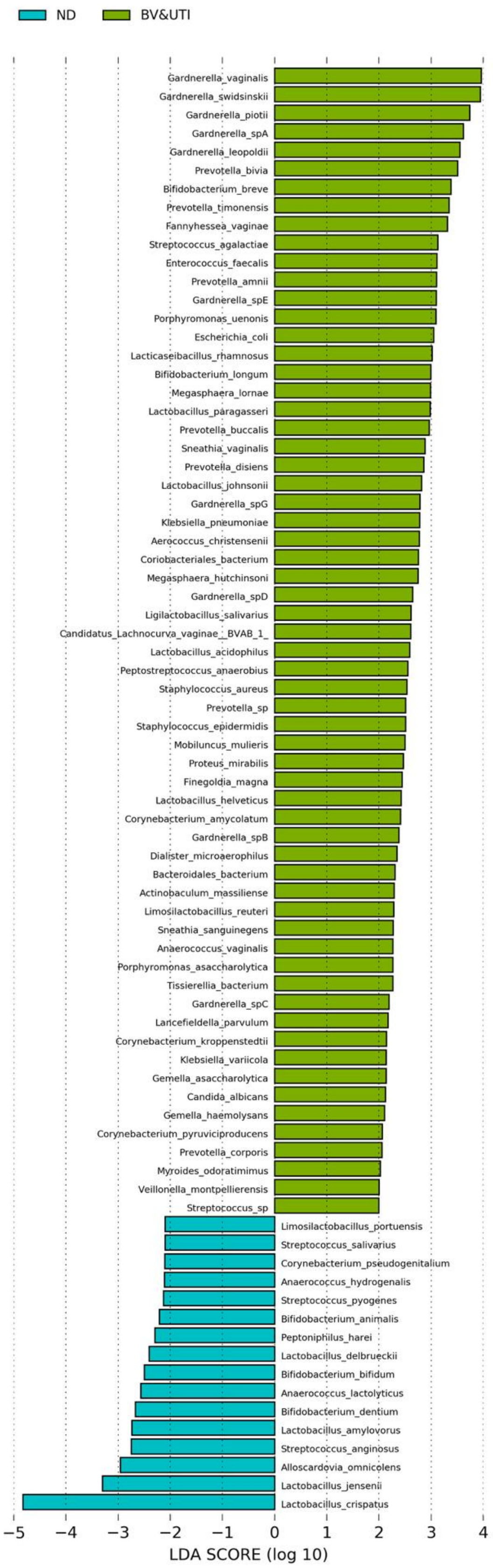
ND
BV&UTI
Gardnerella_vaginalis
Gardnerella_swidsinskii
Gardnerella_piotii
Gardnerella_spA
Gardnerella_leopoldii
Prevotella_bivia
Bifidobacterium_breve
Prevotella_timonensis
Fannyhessea_vaginae
Streptococcus_agalactiae
Enterococcus_faecalis
Prevotella_amnii
Gardnerella_spE
Porphyromonas_uenonis
Escherichia_coli
Lacticaseibacillus_rhamnosus
Bifidobacterium_longum
Megasphaera_lornae
Lactobacillus_paragasseri
Prevotella_buccalis
Sneathia_vaginalis
Prevotella_disiens
Lactobacillus_johnsonii
Gardnerella_spG
Klebsiella_pneumoniae
Aerococcus_christensenii
Coriobacteriales_bacterium
Megasphaera_hutchinsoni
Gardnerella_spD
Ligilactobacillus_salivarius
Candidatus_Lachnocurva_vaginae__BVAB_1_
Lactobacillus_acidophilus
Peptostreptococcus_anaerobius
Staphylococcus_aureus
Prevotella_sp
Staphylococcus_epidermidis
Mobiluncus_mulieris
Proteus_mirabilis
Finegoldia_magna
Lactobacillus_helveticus
Corynebacterium_amycolatum
Gardnerella_spB
Dialister_microaerophilus
Bacteroidales_bacterium
Actinobaculum_massiliense
Limosilactobacillus_reuteri
Sneathia_sanguinegens
Anaerococcus_vaginalis
Porphyromonas_asaccharolytica
Tissierellia_bacterium
Gardnerella_spC
Lancefieldella_parvulum
Corynebacterium_kroppenstedtii
Klebsiella_variicola
Gemella_asaccharolytica
Candida_albicans
Gemella_haemolysans
Corynebacterium_pyruviciproducens
Prevotella_corporis
Myroides_odoratimimus
Veillonella_montpellierensis
Streptococcus_sp
Limosilactobacillus_portuensis
Streptococcus_salivarius
Corynebacterium_pseudogenitalium
Anaerococcus_hydrogenalis
Streptococcus_pyogenes
Bifidobacterium_animalis
Peptoniphilus_harei
Lactobacillus_delbrueckii
Bifidobacterium_bifidum
Anaerococcus_lactolyticus
Bifidobacterium_dentium
Lactobacillus_amylovorus
Streptococcus_anginosus
Alloscardovia_omnicolens
Lactobacillus_jensenii
Lactobacillus_crispatus
−5 −4 −3 −2 −1 0 1 2 3 4
LDA SCORE (log 10)

C)

**D)**

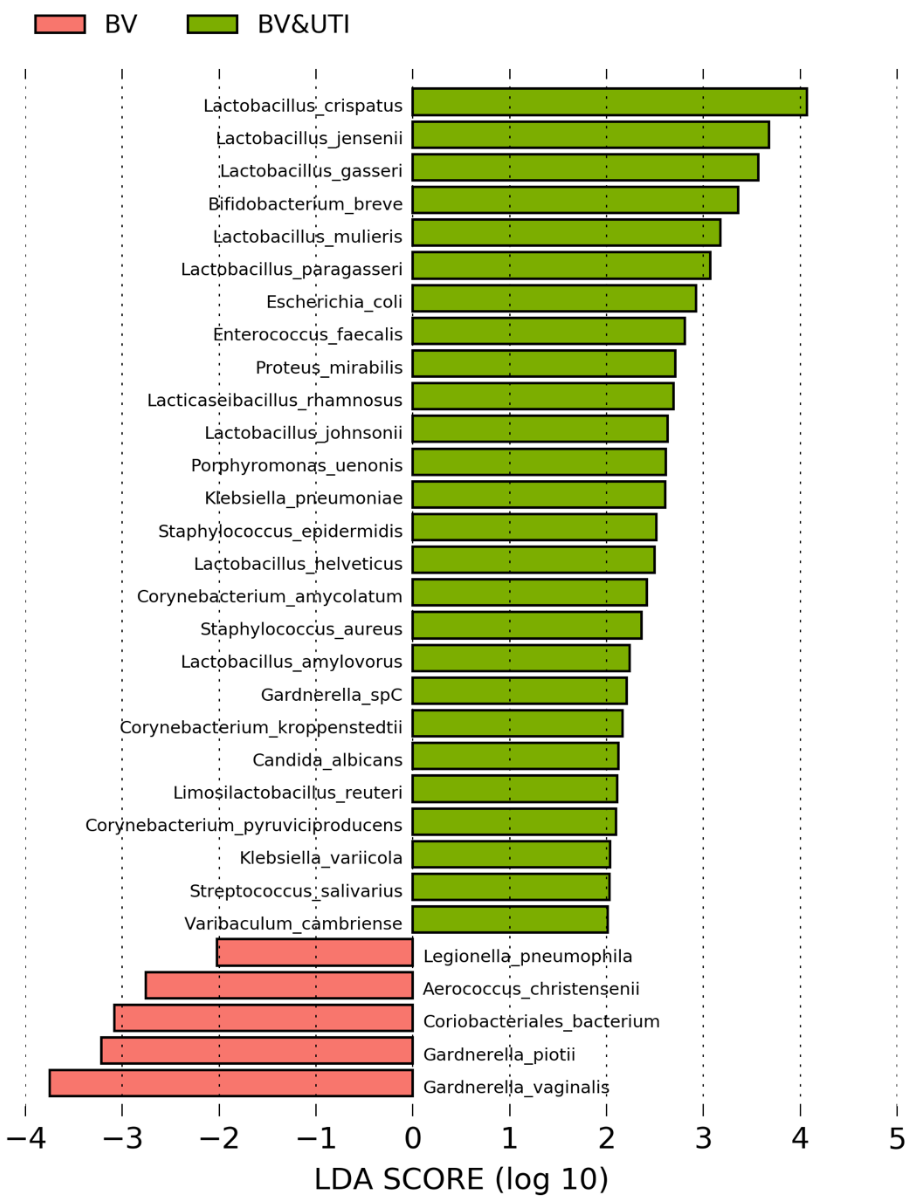


**Supplementary Figure 2:** LDA Scores. Species enriched in the pairwise comparisons with an LDA score > 2.0 are displayed for **(A)** ND vs UTI, **(B)** ND vs BV, **(C)** ND vs BV&UTI, **(D)** BV vs BV&UTI.

**Supplementary Table 1**: Comparisons adjusted odds ratio (95% CI) between groups under Category Variables. Race & ethnicity categories were compared to '*White'* women as a reference. '*Not Pregnant'* women were used as reference for the other categories under Pregnancy Status, and women '*Not in menopause'* were used as a reference for the other categories under Menopause Status. Pairwise comparisons of adjusted least-squares means differences for the Numerical Variables are shown. Bonferroni adjustments were performed for multiple comparisons among the subgroups.

**Supplementary Table 2**: Comparison adjusted odds ratio and 95% confidence interval (CI) on **(A)** CST (Community State Types) distributions, **(B)** VALENCIA Type distributions, and **(C)** CST distributions between clinically diagnosed <YE> vs undiagnosed/suspected <TH> infections. *P*-values * <0.05, ** <0.01, *** <0.001, **** <0.0001 by overall Chi-square test. Significant differences were noted in **bold** and colored.

**Supplementary Table 3**: Symptom severity was quantitatively evaluated at sampling timepoint using a 12-parameter symptom questionnaire that includes the symptoms listed. Participants rated each symptom on a scale: MI = mild, MO = moderate, SE = severe. Participants experiencing moderate to severe symptoms were categorized in the *symptoms* group, while participants with mild or no symptoms were grouped in the *no-symptoms* category and used as reference. Adjusted odds ratio and 95% CI between study group pairwise comparisons were determined. Significant differences were noted in colors.

**Supplementary Table 4**: Comparison abundance of uropathogens based on clinically diagnosed <YE> and undiagnosed/suspected <TH> infection, or both <ALL> using adjusted LS-mean and Bonferroni adjustment for multiple testing. *P*-values * <0.05, ** <0.01, *** <0.001, **** <0.0001. Significant differences were noted in bold and colored.

**Supplementary Table 5**: Comparison adjusted odds ratio (CI 95%) of different treatment types used within the past 30 days from sample submission. *P*-values * <0.05, ** <0.01, *** <0.001, **** <0.0001 by Pearson's Chi-square test. Significant differences were noted in **bold.**

**Supplementary Table 6**: Comparison abundance of uropathogens based on whether **(A)** they took antibiotics (+) or not (-), or both (ALL) using adjusted LS-mean and Bonferroni adjustment for multiple testing. Antibiotics were used no later than 7 days before sample submission. **(B)** We divided the comparisons between clinically diagnosed <YE> and suspected <TH> infection. *P*-values * <0.05, ** <0.01, *** <0.001, **** <0.0001. Significant differences were noted in bold and colored.

**Supplementary Table 7**: Comparison abundance of *Gardnerella* species based on whether they took antibiotics (+) or not (-), or both (ALL) using adjusted LS-mean and Bonferroni adjustment for multiple testing. *P*-values * <0.05, ** <0.01, *** <0.001, **** <0.0001. Antibiotics were used no later than 7 days before sample submission. Significant differences were noted in **bold.**
